# Impacts of climate change on high priority fruit fly species in Australia

**DOI:** 10.1101/567321

**Authors:** Sabira Sultana, John B. Baumgartner, Bernard C. Dominiak, Jane E. Royer, Linda J. Beaumont

## Abstract

Tephritid fruit flies are among the most destructive horticultural pests and pose risks to Australia’s multi-billion-dollar horticulture industry. Currently, there are 11 pest fruit fly species of economic concern present in various regions of Australia. Of these, nine are native to this continent (*Bactrocera aquilonis, B. bryoniae, B. halfordiae, B. jarvisi, B. kraussi, B. musae, B. neohumeralis, B. tryoni* and *Zeugodacus cucumis*), while *B. frauenfeldi* and *Ceratitis capitata* are introduced. To varying degrees these species are costly to Australia’s horticulture through in-farm management, monitoring to demonstrate pest freedom, quarantine and trade restrictions, and crop losses. Here, we used a common species distribution modelling approach, Maxent, to assess habitat suitability for these 11 species under current and future climate scenarios. These projections indicate that the Wet Tropics is likely to be vulnerable to all 11 species. The east coast of Australia will likely remain vulnerable to multiple species until at least 2070. Both the Cape York Peninsula and Northern Territory are also likely to be vulnerable, however, extrapolation to novel climates in these areas decrease confidence in model projections. The climate suitability of current major horticulture regions in north-western Australia, the Northern Territory, southern-central regions of New South Wales and southern Victoria to these pests is projected to increase as climate changes. Our study highlights areas at risk of pest range expansion in the future, to guide Australia’s horticulture industry in developing effective monitoring and management strategies.

## Introduction

Tephritid fruit flies are one of the most destructive and economically significant pest insect families, attacking a wide range of fruit and vegetables. While the family contains more than 4000 species, around 350 are recognized as economically important horticultural pests [1] that have significant impacts on global horticultural production and market access. In Australia, the average annual value of crops susceptible to fruit flies is ∼$4.8 billion [1], and the National Fruit Fly Strategy has identified 46 species as ‘high priority pests’ [2] of concern. The majority of these species are exotic to Australia, primarily found in South-East Asia and the South Pacific [1, 2], and are yet to establish populations in Australia. Of the 11 species that are currently present in Australia [1-3] (Table 1), seven are reported to cause significant economic losses (*Bactrocera aquilonis, B. jarvisi, B. neohumeralis, B. musae, B. tryoni, Ceratitis capitata*, and *Zeugodacus cucumis*) [1, 4]. Combined, these species infest a wide variety of hosts, with some (e.g. *B. frauenfeldi, B. jarvisi, B. neohumeralis, B. tryoni* and *Ceratitis capitata*) [3] being highly polyphagous.

**Table 1.** Eleven economically-significant tephritid pest species present in Australia. Eleven tephritid pest species present in Australia, including nine natives (*B. aquilonis, B. bryoniae, B. halfordiae, B. jarvisi, B. kraussi, B. musae, B. neohumeralis, B. tryoni* and *Z. cucumis*) and two introduced species (*B. frauenfeldi* and *C. capitata*), and their major commercial hosts.

| Species | Common name | Geographical Range | Major Commercial Hosts [32] | References |
| --- | --- | --- | --- | --- |
| *Bactrocera aquilonis* (May) | Northern Territory fruit fly | Top End of the Northern Territory (NT), northern areas of Western Australia | Bell pepper, tomato, lemon, mandarin, grapefruit, apple, mango, peach | 1 |
| *Bactrocera bryoniae* (Tryon) | N/A | Torres Strait Islands, mainland Queensland, northern Western Australia, NT, NSW as far south as Sydney | Chilli, tomato | 3 |
| *Bactrocera halfordiae* (Tryon) | Halfordia fruit fly | North Queensland south to the Sydney region in NSW | Citrus | 3 |
| *Bactrocera jarvisi* (Tryon) | Jarvis' fruit fly | North-western Western Australia, NT, north-west Queensland, eastern Australia from Cape York to Sydney, NSW | Mango, peach, banana, pear, apple, pawpaw, persimmon | 3 |
| *Bactrocera kraussi* (Hardy) | Krauss' fruit fly | Torres Strait Islands, northeast Queensland as far south as Townsville | Grapefruit, mandarin, orange, mango, peach and banana | 3, 7 |
| *Bactrocera musae* (Tryon) | Banana fruit fly | Torres Strait Islands, northeast Queensland as far south as Townsville | Banana | 3, 7 |
| *Bactrocera neohumeralis* (Tryon) | Lesser Queensland fruit fly | Torres Strait Islands, eastern Queensland, northern NSW | Mango, papaw, persimmon, avocado, banana, passionfruit, apple, apricot, plum, peach, citrus, capsicum, chilli, tomato | 3 |
| *Bactrocera tryoni*<br>(Froggatt) | Queensland fruit<br>fly (Qfly) | Central and Top End of NT, eastern<br>Australia, Victoria | Mango, papaw, avocado,<br>grapefruit, passionfruit,<br>strawberry, peach, pear, apple,<br>banana, persimmon, chilli,<br>capsicum, tomato, eggplant | 3,31 |
| *Zeugodacus cucumis*<br>(French) (formerly<br>*Bactrocera cucumis*) | Cucumber fruit<br>fly | Eastern Queensland, north-eastern<br>NSW, NT | Cucumber, pumpkin, zucchini,<br>squash, passionfruit, tomato,<br>pawpaw | 3 |
| *Bactrocera*<br>*frauenfeldi* (Schiner) | Mango fruit fly | Native to Papua New Guinea and<br>surrounding islands, spread to<br>Torres Strait Islands and northern<br>Queensland as far south as<br>Townsville | Mango, banana, passionfruit,<br>citrus, chilli | 7 |
| *Ceratitis capitata*<br>(Wiedemann) | Mediterranean<br>fruit fly<br>(Medfly) | Native to Africa, spread to the<br>Mediterranean regions, Western<br>Australia, occasional detections in<br>South Australia and NT are<br>eradicated. | Mango, papaw, apple, peach,<br>pear, citrus | 3, 28, 31 |

The distributions of Australia’s pest fruit fly species are influenced by their climatic tolerances and the distributions of their hosts. *Bactrocera* originated in tropical regions, and have their highest richness in rainforests [5]. However, over the last 100 years, as horticulture has proliferated across Australia, some species have expanded their geographic range and host breadth [6]. Of the 11 high priority fruit fly species presently on the continent, three are currently restricted to north-east Queensland (*B. frauenfeldi, B. kraussi* and *B. musae*) [7]. In contrast, the geographic range of *B. neohumeralis* (Lesser Queensland fruit fly) extends along eastern Australia, from Queensland to central New South Wales (NSW) [3, 4, 7]. Previous climatic analysis indicates that this species also has the potential to establish elsewhere in northern Australia [4]. The remaining species have substantially wider climate tolerances, and are found across broad regions of the continent. For instance, *B. tryoni* (Qfly) ranges across much of eastern Australia, eastern Queensland and northern regions of the Northern Territory [8]. *Bactrocera jarvisi* (Jarvis’ fruit fly) extends from northwest Western Australia, across the Northern Territory to northern Queensland and the Torres Strait Islands [4, 9], and, in favourable years, may spread down the east coast of Australia into northern coastal NSW [4, 9]. Hence, *B. jarvisi* and *B. tryoni* have overlapping geographic ranges and infest many of the same hosts [4]. *Ceratitis capitata* (Medfly) originated from the Afrotropical region [10], and was introduced into the Perth area (Western Australia) in the late 1800s [4]. Before quarantine controls were developed, this species spread to NSW, Victoria, and other parts of Australia [11]. However, for reasons that remain unclear, Qfly is believed to have displaced Medfly throughout most of its former Australian range [12], and now Medfly is confined to Western Australia, with occasional detections in South Australia [13].

Under current climate conditions, most of these 11 fruit fly species pose threats to Australia’s horticulture industries, as well as to backyard growers. As such, controlling fruit flies is imperative for the viability of Australian horticulture, necessitating in-farm management and pest treatment, monitoring to demonstrate pest freedom, and quarantine and trade restrictions [1, 2]. These controls, along with loss of market access, are estimated to cost Australian growers $100 million per annum [4], in addition to losses of up to $159 million per annum due to infestation of fruit and vegetable crops [14].

For those areas where fruit flies are found, the annual cost, as reported in 2012, of bait and cover spray, as well as post-harvest treatments, amount to $269 ha^−1^ and $62.36 tonne^−1^, respectively [15], while maintaining fruit fly free areas is estimated to exceed $28 million per annum based on data from 2009-2011 [16]. However, restrictions were recently placed on the use of insecticides to control fruit flies due to concerns about toxicity [17], with dimethoate and fenthion suspended or highly restricted for many horticultural crops [17-20]. Other approaches, including Sterile Insect Techniques, are now being explored. Regardless, it has been estimated that the annual likelihood of an incursion by an exotic fruit fly species is 21% [15], and the annual cost of eradicating these incursions is ∼$13 million [16], with rapid responses to outbreaks being crucial for eradication [21]. Even brief incursions can result in significant economic damage due to market access restrictions that may be imposed. However, climate change is likely to alter the distribution of suitable habitat for fruit fly species and areas vulnerable to outbreaks, and this could have serious repercussions for Australian horticulture [22].

Previous studies [22-24] have used the semi-mechanistic species distribution model (SDM), CLIMEX, to estimate the potential geographic distributions of several high priority fruit fly species, based on their performance along climatic gradients. While highly useful in furthering our understanding of climate impacts on fruit flies, these studies have either focused on other countries [24-26] or have explored global patterns of the distribution of suitable climate [22, 23]. Here we assess how climate change may result in shifts to the distribution of suitable habitat for the 11 high priority fruit fly species present in Australia, using the correlative SDM, Maxent [27]. This SDM has been used extensively to assess the distribution of suitable habitat for a broad range of pests and invasive species [26, 28-31]. We also highlight areas at risk of pest range expansion, to guide Australia’s horticulture industries in development of effective monitoring and management strategies.

## Methodology

### Species occurrence data

We collected occurrence data for the 11 species from five main sources: the Australian Plant Pest Database (APPD; http://www.planthealthaustralia.com.au/resources/australian-plant-pest-database, accessed 15th March 2017), the Atlas of Living Australia (ALA; http://www.ala.org.au, 22nd December, 2016), the Global Biodiversity Information Facility (GBIF, https://www.gbif.org, 28th June, 2017), trap data, and existing literature. APPD is a national digital database of plant pest and pathogen specimens held within herbaria and insect collections across Australia. It is a powerful tool for market access and emergency responses to pest incursion, and supports associated research activities. ALA is Australia’s largest digital database of species occurrence records, containing information from a wide array of data providers including Australia’s major museums and government departments. GBIF provides similar data at a global scale. Before downloading data from APPD, ALA and GBIF, we applied filters to restrict records to those that were resolved to species-level, were dated no earlier than 1 January 1950, contained valid geographic coordinates, and were not flagged as ‘environmental outliers’.

We also collected trap data from various state government departments (Biosecurity and Food Safety, Department of Primary Industries, NSW; Biosecurity Queensland and the Queensland Department of Agriculture and Fisheries; Department of Economic Development, Jobs, Transport and Resources, Victoria; and Department of Primary Industries and Regions South Australia (PIRSA)). Trap data from these sources were collected at different periods from 1996 to 2017. Finally, we also obtained occurrence data from the literature [1-4, 6, 7, 11, 33-37].

### Major commercial fruit and vegetable hosts

For each of the 11 fruit fly species, we compiled information on the major commercial hosts on which infestation has been recorded. For this purpose, we defined major fruit and vegetable host species according to the Australian Horticulture Statistics Handbook (HSHB; www.horticulture.com.au) for the year 2016/2017 [32]. This document consolidates horticulture statistics of interest to industry members and other stakeholders. The data contained in HSHB were derived from the Australian Bureau of Statistics, projects funded by Hort Innovation, international trade sources and horticulture industry representative bodies where available.

### Climate data

For current and future climate conditions we used the bioclimatic variables available within the WorldClim database, at a spatial resolution of 30 arc-seconds [38] (approximately 1 km; http://www.worldclim.org). These data, based on meteorological records for the period 1960– 1990, comprise 19 climatic variables, 11 of which are temperature-based while eight relate to precipitation. Combined, the data represent annual trends, seasonality, and limiting or extreme environmental conditions. Assuming that host plants are available, temperature and moisture are the key factors influencing fruit fly reproduction and survival [18, 39]. Thus, these variables were chosen as predictor candidates based on the fruit flies’ biology and ecological requirements, and similar habitat suitability studies undertaken on other insects [40]. For each species, we identified a set of ecologically-relevant variables, with minimal collinearity, that resulted in high predictive power for the model [41] (described below).

When projecting the future suitability of habitat, we considered a range of climate scenarios to acknowledge this important aspect of uncertainty. We choose six global climate models (GCMs) recommended by CSIRO as being useful for Australian climate impact assessments [42], and for which data were available at our chosen spatial resolution for the Representative Concentration Pathway 8.5 (RCP8.5) [43]. These GCMs included: CanESM2 (The Second Generation of Canadian Earth System Model); ACCESS1.0 (The Australian Community Climate and Earth System Simulator); MIROC5 (Model for Interdisciplinary Research on Climate); HadGEM2-CC (Hadley Centre Global Environmental Model Version 2 Carbon Cycle); NorESM1-M (The Norwegian Earth System Model-Part-1); and GFDL-ESM2M (Global Coupled Climate Carbon Earth System Model Part-1). The CanESM2 model projects a hot future with drying across central regions and higher precipitation in the north-east. The ACCESS1.0 model projects a hot and dry future across most areas of Australia, while MIROC5 projects moderate warming, with drying in the north-east and south-west but higher precipitation in central Australia. NorESM1-M projects moderate warming. HadGEM2-CC and GFDL-ESM2M project a hot future with warming typically in central regions. We downloaded the 19 bioclimatic variables from these six models for 20-year periods centred on 2030, 2050 and 2070 (http://www.climatechangeinaustralia.gov.au). These data were then reprojected to a spatial resolution of 1 × 1 km (Australian Albers Equal Area, EPSG: 3577) via bilinear interpolation, using the gdalwarp function provided by the R package gdalUtils [44] in R version 3.3.3 [45].

### Species Distribution Models

We used the machine learning approach, Maxent (v3.3.3k [27]), to assess habitat suitability for species under current and future climate scenarios. Maxent accommodates presence-only data and has performed well in multimodel assessments [46]. It produces a continuous probability surface, which can be interpreted as an index of habitat suitability given the predictor variables included in model calibration. Detailed descriptions of Maxent are given elsewhere [47, 48]. We optimized models by assessing the effects of different combinations of feature types, of competing predictor sets deemed ecologically sensible *a priori*, and of the extent of regularization on model performance. We found that Maxent performed best when product (first-order interactions), linear and quadratic features were used, with a regularization multiplier of 1 (the default), and used this configuration to calibrate our final models.

Maxent requires background data, to which it compares the environmental characteristics of presence locations. There is flexibility for users to specify which points to use as background, as well as the number of records and the spatial extent from which they are chosen [47]. Following Ihlow *et al* [49], we manually generated background points by randomly selecting 100,000 cells from terrestrial areas within 200 km of occurrence records of the target species. Our choice of background achieves a balance between fine-scale discrimination of suitable and unsuitable sites along environmental gradients, and generalization of model predictions.

To assess model performance, we used five-fold cross-validation to reduce model errors that may occur from the random splitting of data into test and training subsets. The performance of each model was evaluated using the area under the receiver operating characteristic curve (AUC), which describes the consistency with which a model ranks randomly chosen presence sites as more suitable than randomly chosen background sites. AUC ranges from 0 to 1, with a value of 0.50 indicating discrimination ability no better than random, while values greater than 0.75 indicates that the model has a discriminative ability that is better than “fair” [50]. Cross-validated AUC scores were presumed to reflect the performance of a single final model for each species, which used all available data.

Following previous studies of pest species [26], continuous habitat suitability scores projected by Maxent models were converted to binary layers (0 = unsuitable, 1 = suitable) using the 10th percentile training presence threshold (i.e. the value that corresponds to 10% training omission). We note that the selection of a threshold value is subjective and may vary depending upon the goals of the study [51], thus we also provide continuous output for current climate as supplemental data (**S1-11 Figs**). For each species, the six binary suitability grids (i.e., one for each GCM, with cells assigned 0 when unsuitable and 1 when suitable) for each time period were summed to produce a consensus map, identifying agreement about the suitability of grid cells across the six climate scenarios. Each species’ consensus map was then converting to a binary map indicating whether cells were projected to be suitable under the majority of GCMs (i.e., suitable in < 4 GCMs = 0, suitable in 4 or more = 1). The resulting binary maps were summed across species to identify hotspots - grid cells suitable for multiple pest species. Finally, we compared the distribution of hotspots to that of major horticultural crops.

When projecting models, extrapolation to conditions beyond the range of the training data may be unreliable. Following Elith *et al.* [52] we developed MESS (multivariate environmental similarity surface) maps to identify regions of extrapolation [52]. By revealing areas with novel environmental conditions, MESS maps can be used as a projection mask, highlighting regions for which projections are unreliable, or as a quantitative measure of prediction uncertainty [52]. To assess how novel environments may alter projections of habitat suitability, for each species we visually compared maps of habitat suitability that included projections made in areas with novel environmental conditions to maps that classified these areas as unsuitable.

All modelling and post-modelling analyses and calculation of statistics were performed in R version 3.1.2 [53]. We used the sp [54] and raster [55] packages for preparation and manipulation of spatial data, the dismo package to fit Maxent models, and custom R code for rapid projection of fitted models.

## Results

### Model Performance

Model performance for all species was better than random, with average cross-validated AUC ranging from 0.815 (SD = 0.05; *B. frauenfeldi*) to 0.907 (SD = 0.02; *B. neohumeralis*) **(S1 Table)**.

#### Bactrocera aquilonis

Our model suggested that suitable habitat for *B. aquilonis* currently exists in the northern regions of the Northern Territory and Western Australia, as well as in northern Queensland, where this fly has not been reported (**S1A-1B Figs**). The variables with the highest permutation importance were precipitation of the wettest quarter (68.9%) and annual mean temperature (28.9%) **(S1 Table)**.

As the century progresses, the geographic extent of suitable habitat for this species is projected to increase and shift southwards under all six scenarios **(S1C-1E Figs).** Further, most areas currently suitable are projected to remain so until at least 2070. By 2030, 21.5% of Australia (i.e. ∼1,600,100 km^2^) is projected to be climatically suitable under at least one climate scenario **(S3 Table)**, and this is projected to increase to 39.9% (∼3,000,100 km^2^) by 2070 **(S3 Table)**. However, a substantially smaller area is projected suitable under all six scenarios (2030 = 9.2%, ∼700,100 km^2^; 2070 = 20.7%, ∼1,590,800 km^2^) **(S3 Table)**. This includes northern Western Australia, much of the Northern Territory, and north-western Queensland **(S1C-1E Figs).**

Key horticultural crops for *B. aquilonis* are *Mangifera indica* (mango), *Citrus × paradisi* (grapefruit), *Malus domestica* (apple), *Prunus persica* (peach) and *Citrus sp.* (citrus) **(S4 Table)**. The major regions where these crops are currently grown include the Northern Territory and north-east Western Australia. These regions will likely remain suitable for *B. aquilonis* until at least 2070. Similarly, fruit growing regions in the Wet Tropics (north-east Queensland) are likely to increase in suitability in the future. Other major host-plant growing regions in the south and east of the continent will likely remain unsuitable **(S1 Fig).**

#### Bactrocera bryoniae

Current suitable habitat for *B. bryoniae* is projected to occur along the northern and eastern coastlines **(S2A-2B Figs).** Temperature annual range and precipitation of the driest month contributed the most to the model for this species (42.2% and 27.4%, respectively) **(S1 Table)**.

By 2070, suitable habitat is projected to increase under all scenarios except GFDL-ESM2M (which projects a hot, very dry future) **(S2C-2E Figs; S2 Table)**, expanding to the southern coastlines of Victoria and Western Australia. By 2030, 13.5% of Australia (i.e. ∼1,038,000 km^2^) is projected to be suitable for *B. bryoniae* under at least one of the climate scenarios, increasing to 20.6% (∼1,500,700 km^2^) by 2070 **(S3 Table)**. Under 1-3 scenarios, suitable habitat is projected to shift inland in Queensland and NSW. However, the amount of habitat projected to remain suitable under all six scenarios remains relatively stable from 2030-2070 (i.e. spanning 6.5–7.1% of the continent) **(S3 Table)**.

The major horticultural host for *B. bryoniae* is *Capsicum annuum* (chilli) **(S4 Table)**. Our model indicates that key growing regions for this crop in Queensland currently contain suitable habitat for *B. bryoniae*, and this will continue to be the case until at least 2070 **(S2 Fig).**

#### Bactrocera frauenfeldi

Currently, suitable habitat for this species is projected to be mostly confined to Cape York Peninsula and the Wet Tropics, although there are also small areas in northern Western Australia and the Northern Territory that are classified as suitable, but from which the species has not been recorded **(S3A-3B Figs).** The most important variable in the model for *B. frauenfeldi* was precipitation of the wettest quarter (75.4%) **(S1 Table)**.

As the century progresses, suitable habitat is projected to increase under all scenarios except CanESM2 **(S2 Table)**. This scenario projects a hot, very dry future, leading to loss of suitable habitat in northern Queensland by 2050. In contrast, under the ACCESS1.0 scenario (a hot, dry scenario) total range size may increase by 43.6% by 2030. However, by 2070, the Cape York Peninsula region is projected to become unsuitable **(S2 Table)**. Overall, 3.3% of Australia (i.e. ∼250,400 km^2^) is projected to be suitable for *B. frauenfeldi* under at least one scenario by 2030, increasingly slightly to 4% (∼300,000 km^2^) by 2070 **(S3 Table)**. Only 1.3% (∼100,000 km^2^) of the continent is projected to be suitable under all six scenarios by 2070 **(S3 Table)**.

The major crops for *B. frauenfeldi* are *Mangifera indica* (mango) and *Carcica papaya* (pawpaw) **(S4 Table)**. Major production regions in north-western Northern Territory may remain suitable for this species until at least 2070, although there is substantial uncertainty across the climate scenarios. In contrast, it is very likely that the Wet Tropics will remain suitable until at least 2070, irrespective of the climate scenario **(S3 Fig).**

#### Bactrocera halfordiae

Suitable habitat for *B. halfordiae* is currently found in the Wet Tropics and subtropics from north Queensland to eastern New South Wales **(S4A-4B Figs).** Precipitation of the driest month (66.8%) and annual mean temperature (32.3%) contributed most to this model **(S1 Table)**.

The geographic extent of suitable habitat is projected to vary considerably across the six climate scenarios, however as the century progresses, gains in new habitat may exceed losses under some scenarios (e.g. see ACCESS and MIROC5 in **S2 Table**). In contrast, under the CanESM2 scenario (which projects a hot future, drying across central regions and higher precipitation in the north-east), total range size may decline by 65.1% by 2030 **(S2 Table)**, mostly due to contractions in the south and east, although limited gains in habitat may occur in northern Australia. Similarly, under the HadGEM2 scenario (which projects substantial drying across some regions of the continent), 51.4% of current suitable habitat is projected to be lost by 2030, although by 2070, range expansions may exceed losses **(S2 Table)**. By 2030, 6.9% of Australia (i.e. ∼500,800 km^2^) is projected to be suitable for *B. halfordiae* under at least one climate scenario, increasing to 9.9% (∼750,900 km^2^) by 2070 **(S3 Table)**.

Comparing the current and future distribution of suitable habitat for this fly with the major growing regions for its host plants indicates that crops in the Wet Tropics may continue to be at risk, until at least 2070. However, only 1–2 scenarios project horticultural regions in southern Queensland to retain suitable climate **(S4C-4E Figs).** Although horticultural regions along the NSW-Victorian border are currently unsuitable for *B. halfordiae*, some models project these areas to become suitable between 2050–2070 **(S4 Fig).**

#### Bactrocera jarvisi

Current suitable habitat for this species is projected to be mostly confined to northern Western Australia, the Top End of the Northern Territory, and eastern Australia from Cape York to NSW **(S5A-5B Figs).** Annual mean temperature (38.0%) and precipitation of driest month (37.2%) had the highest contributions to the model for this species **(S1 Table)**. There is substantial consensus across the six scenarios that regions currently suitable for *B. jarvisi* will remain so until at least 2070 **(S5C-5E Figs; S2 Table).** In addition, across some models, gains are projected to occur in Western Australia, the Northern Territory and central Queensland. For instance, under the CanESM2 scenario, total range size may increase by 45.5% to 58.7% between 2030 and 2070. Similarly, under the hot, very dry scenario projected by ACCESS1.0, 62.5% of future suitable habitat may occur in new areas by 2070 **(S2 Table)**. By 2030, 25.8% of Australia (i.e.∼1,980,000 km^2^) is projected to be suitable for *B. jarvisi* under at least one of the climate scenarios, and due to subsequent gains in suitable habitat, this may increase to 44.3% (∼3,390,400 km^2^) by 2070 **(S3 Table)**.

Comparing the distribution of suitable habitat for this fly with that of its major host crops indicates that crops currently grown in the Top End of the Northern Territory, and in eastern Australia from Cape York to New South Wales, may continue to be at risk until at least 2070. Other major host-plant growing regions in the south and west of the continent will also remain suitable for this species until 2070 **(S5 Fig).**

#### Bactrocera kraussi

Suitable habitat for *B. kraussi* is projected to occur across the northern tip of Australia and northeast Queensland, as far south as Townsville **(S6A-6B Figs).** Precipitation of the wettest quarter (75.19%) had the highest contribution to the model of *B. kraussi* **(S1 Table)**.

The geographic extent of suitable habitat is projected to increase under all six scenarios **(S6C-6E Figs; S2 Table).** For example, under the NorESM1-M scenario (a moderate warming scenario with little precipitation change), an area equivalent to 29.6% of current suitable habitat is projected to be gained by 2070 **(S2 Table)**. By 2030, 3.1% of Australia (i.e.∼240,600 km^2^) is projected to be suitable for *B. kraussi* under one or more of the climate scenarios, increasing to 3.8% (∼280,300 km^2^) by 2070 **(S3 Table)**. This includes north-east Queensland **(S6C-6E Figs).** Horticultural production regions in northeast Queensland as far south as Townsville may remain suitable for this species by 2070, although production regions in the south are likely to remain unsuitable **(S6 Fig)**.

#### Bactrocera musae

Current suitable habitat for *B. musae* is predicted from the Torres Strait Islands through to the Wet Tropics **(S7A-7B Figs).** The most important variable in the model for *B. musae* was precipitation of the wettest quarter (78.7%) **(S1 Table)**.

Projections for this species under climate change are similar to those for *B. kraussi*. Under the CanESM2 scenario, 22.2% of current suitable habitat is projected to be lost by 2030, although by 2070, range expansions are projected to exceed losses **(S2 Table)**. However, in NorESM1-M (a moderate warming scenario with little precipitation change), total range size may increase by ∼43.5% by 2070 **(S2 Table)**. By 2030, 3.4% of Australia (i.e.∼260,100 km^2^) is projected to be suitable for *B. musae* under at least one of the climate scenarios **(S3 Table)**, increasing to 4.2% (∼300,600 km^2^) by 2070 **(S3 Table)**.

*B. musae* mainly attacks *Musa × paradisiaca* (banana), the production areas for which are located primarily in tropical and subtropical regions of the continent **(S4 Table)**. The major commercial growing region in the Wet Tropics is projected to remain climatically suitable for this species until at least 2070 (S8 Fig).

#### Bactrocera neohumeralis

Current suitable habitat for this species is projected to be mostly confined to the Torres Strait Islands, eastern Queensland, and north eastern NSW south to Wollongong **(S8A-8B Figs).** Precipitation of the wettest month (47.4%) contributed most to the model for *B. neohumeralis* **(S1 Table)**.

As the century progresses, considerable differences in suitable habitat are projected across the six scenarios. For example, under the CanESM2 scenario, 24.2% of current suitable habitat is projected to be lost by 2030, although by 2070, range expansions are projected to exceed losses **(S2 Table)**. Similarly, under the hot, very dry scenario simulated by GFDL-ESM2M, total range size may decline by 32.4% by 2030, mostly due to contractions in the south and east, although limited gains in habitat may occur in northern Australia **(S8C-8E Figs; S2 Table).** By 2030, ∼6.9% of Australia (i.e.∼500,500 km^2^) is projected to be suitable for *B. neohumeralis* under at least one of the climate scenarios, increasing to 14% (∼1,000,100 km^2^) by 2070 **(S3 Table)**.

Production regions in eastern Queensland and north-eastern NSW will likely remain suitable for this species until at least 2070, although there is substantial uncertainty across the climate scenarios. In contrast, regions along the NSW-Victorian border and further south are projected to remain unsuitable for *B. neohumeralis* **(S8 Fig).**

#### Bactrocera tryoni

Highly suitable habitat for *B. tryoni* is projected to occur along south-western Western Australia, south-eastern South Australia, Victoria, and eastern Australia from Cape York to NSW **(S9A-9B Figs).** Coastal zones in northern Western Australia, the Northern Territory and the eastern half of Tasmania have moderate suitability **(S9A-9B Figs).** Annual mean temperature (33.06%) and mean temperature of the coldest month (32.42%) had the highest contributions to the model for this species **(S1 Table)**.

The geographic extent of suitable habitat varies across the six climate scenarios. However, as the century progresses, gains in new habitat may exceed losses under some scenarios (e.g. see ACCESS1.0, MIROC5 and NorESM1-M; **S2 Table).** In contrast, under the GFDL-ESM2M scenario, total range size may decline by 47.9% by 2030 **(S2 Table)**, mostly due to contractions in the south and east. Under the ACCESS1.0 scenario, 26.1% of current suitable habitat is projected to be lost by 2030, although by 2070, range expansions are projected for some regions **(S2 Table)**. By 2030, 26.1% of Australia (i.e.∼1,990,800 km^2^) is projected to be suitable for *B. tryoni* under at least one scenario, increasing to 39.6% (∼3,000,500 km^2^) by 2070 **(S3 Table)**. This includes south-west Western Australia, most of Victoria and eastern Tasmania, eastern Queensland and the northern reaches of the continent **(S9C-9E Figs).**

Comparing the distribution of suitable habitat for this fly with that of its major host crops indicates that key regions for these crops in the Top End of Northern Territory, eastern Australia from Cape York to NSW, Victoria, and some parts of Tasmania, may remain suitable for *B. tryoni* until at least 2070. Major host-plant growing regions in South Australia may also remain suitable for this species until 2070 **(S9 Fig).**

#### Ceratitis capitata

Our model suggests that suitable habitat for *C. capitata* exists throughout Western Australia, the Northern Territory, the east coast of Queensland to NSW and South Australia **(S10A-10B Figs).** Annual mean temperature (47.2%) and mean temperature of the coldest month (46.2%) contributed most to the model for this species **(S1 Table)**.

Under the future climate scenarios, the geographic extent of suitable habitat is projected to increase and shift south **(S10C-10E Figs).** By 2030, 28.3% of Australia (i.e. ∼2,100,400 km^2^) is projected to be suitable for *C. capitata* under one or more scenarios **(S3 Table)**, increasing to 47.1% (∼3,600,800 km^2^) by 2070 **(S3 Table)**. However, under all six scenarios, 13.9% (2030; ∼1,000,000 km^2^) to 17.3% (2070; ∼1,300,000 km^2^) of Australia is likely to be suitable for *C. capitata* **(S3 Table)**. Climate suitability within production regions in the south and west of the continent may gradually increase for this species from 2030 to 2070 **(S10 Fig).**

#### Zeugodacus cucumis

Suitable habitat for *Z. cucumis* is projected to occur along the northern region of Western Australia and the Northern Territory, north-east Queensland, and south along the east coast to NSW **(S11A-11B Figs).** Precipitation of the driest quarter (54.3%) and mean temperature of the coldest quarter (36.2%) had the highest permutation importance in the model for this species **(S1 Table)**.

Under future climate scenarios, the geographic extent of suitable habitat is projected to increase, expanding southward and inland, with most areas that are currently suitable projected to remain so until at least 2070 **(S11C-11E Figs).** However, there is considerable variation among projections for inland regions, indicating higher uncertainty about the future suitability of these regions. For instance, under a scenario where most regions, except central Australia, are projected to become drier (ACCESS1.0), total range size may increase by 68.2% by 2070 **(S2 Table)**, with areas of expansion projected to occur in northern Western Australia, central Australia, eastern Queensland and NSW **(S11C-11E Figs).** Under a warmer/wetter scenario (MIROC5), a substantial westward range expansion is projected in eastern Australia **(S2 Table)**. Hence, although 35.4% (∼2,700,900 km^2^) of Australia is projected to be suitable for this species under at least one scenario for 2070, a much smaller extent is suitable under all six scenarios (10.3%, ∼789,900 km^2^) **(S3 Table)**.

Major commercial growing regions for host crops in Queensland and the Northern Territory are projected to remain climatically suitable for this species until at least 2070 **(S11 Fig).** Other major host-plant growing regions in the south and west of the continent will likely remain unsuitable under the time periods considered in this study **(S11F Fig).**

### Future hotspots of pest fruit flies

For each time period, we stacked habitat suitability maps for all species, to identify regions most likely to contain suitable climate conditions for multiple pest species (i.e. hotspots). As the century progresses, the geographic extent of suitable habitat for most of the 11 species is projected to expand and shift south regardless of whether novel environments are included or excluded **(Fig 1, S12 Fig, Table 2 and S5 Table)**. When regions containing novel climate are included, 31.6% of Australia (i.e. ∼2,400,800 km^2^) is projected to be currently suitable for at least one of the 11 species. By 2070, this may increase to 52.9% (∼4,000,800 km^2^) **(Table 2).** only a small region within Queensland’s Wet Tropics (5,621km^2^) is projected suitable for all 11 species.

**Table 2.** Area (km^2^) and percentage (%) of Australia projected to be suitable for the 11 fruit fly species considered in this study, under current and future climates. Each row of the table indicates the area (and percentage) projected to be suitable now, in 2030, 2050, and 2070, for *n* species, where *n* is given in the “Count” column. Thus, the first row (with Count = 0) gives the area projected to be unsuitable for all 11 species, the row with Count = 1 gives the area projected to be suitable for any one of the 11 species, and the row with Count = 11 gives the area projected to be suitable for all 11 species.

| Count | Suitable area km<sup>2</sup> (% of Australia) |  |  |  |
| --- | --- | --- | --- | --- |
|  | Current | 2030 | 2050 | 2070 |
| 0 | 5,250,277 (68.4%) | 5,175,668 (67.5%) | 4,567,601 (59.5%) | 3,607,211 (47.0%) |
| 1 | 847,090 (11.0%) | 738,118 (9.6%) | 887,083 (11.6%) | 1,122,410 (14.6%) |
| 2 | 850,401 (11.1%) | 806,220 (10.5%) | 1,133,009 (14.8%) | 1,343,740 (17.5%) |
| 3 | 176,163 (2.3%) | 257,558 (3.4%) | 228,976 (2.9%) | 501,960 (6.5%) |
| 4 | 93,709 (1.2%) | 148,806 (1.9%) | 248,057 (3.2%) | 365,032 (4.8%) |
| 5 | 121,923 (1.6%) | 196,041 (2.6%) | 225,006 (2.9%) | 299,590 (3.9%) |
| 6 | 89,766 (1.2%) | 144,056 (1.9%) | 141,026 (1.8%) | 176,864 (2.3%) |
| 7 | 101,763 (1.3%) | 46,306 (0.6%) | 78,803 (1.0%) | 89,042 (1.2%) |
| 8 | 32,385 (0.4%) | 30,536 (0.4%) | 37,543 (0.5%) | 43,472 (0.6%) |
| 9 | 57,695 (0.8%) | 102,927 (1.3%) | 109,316 (1.4%) | 10,8250 (1.4%) |
| 10 | 48,537 (0.6%) | 21,554 (0.3%) | 10,807 (0.1%) | 9,888 (0.1%) |
| 11 | 3,371 (0.0%) | 5,290 (0.1%) | 5,853 (0.1%) | 5,621 (0.1%) |

**Fig 1.**
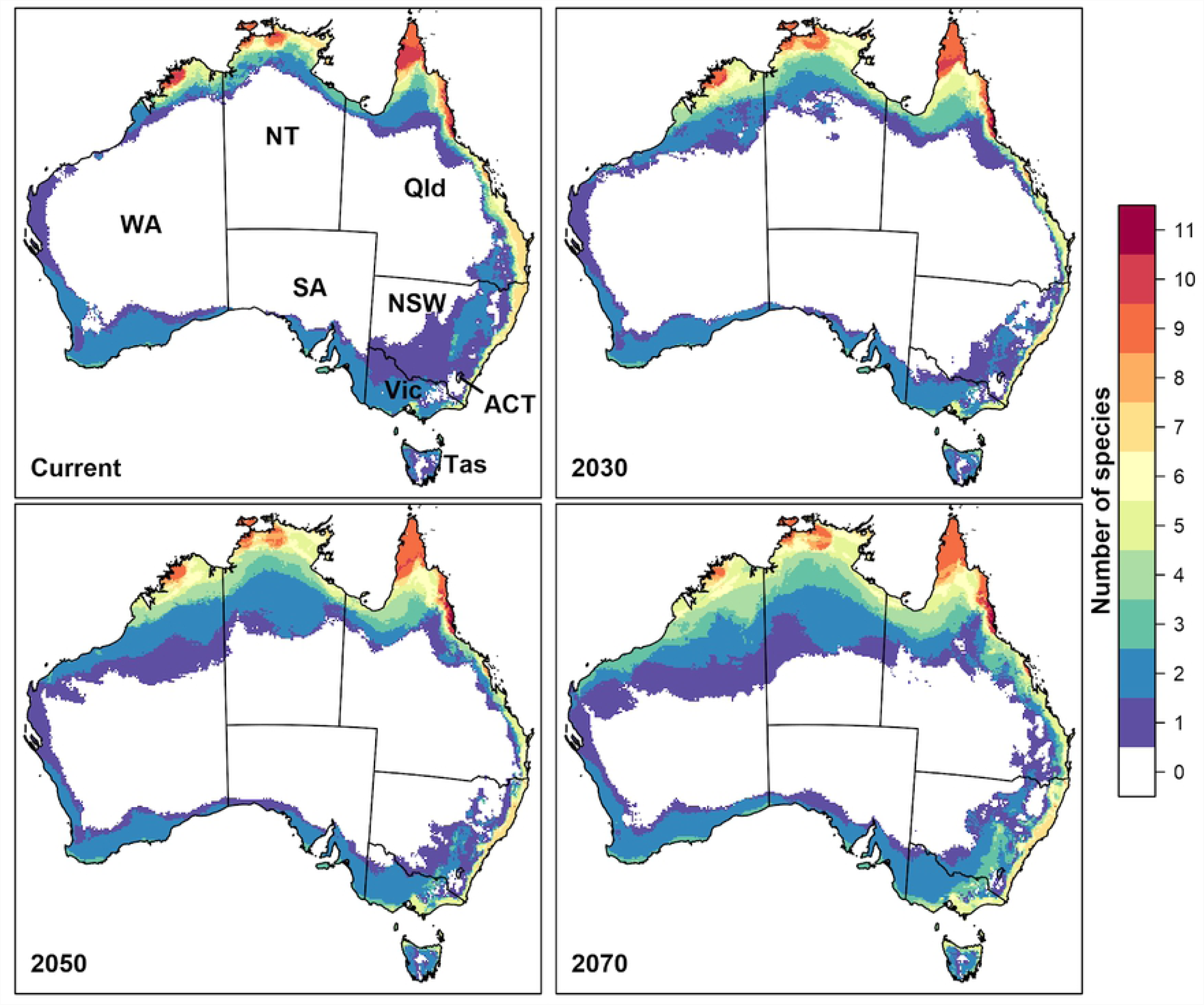
Hotspot maps of habitat suitability for the 11 fruit fly species under climate change, when novel environments are included. Hotspot maps of current and future habitat suitability for 11 fruit flies. Suitability was modelled with Maxent, and thresholded using the 10th percentile of suitability at training presence localities. These maps include projections under novel environments. Colours indicate the number of species for which habitat is projected to be suitable under the majority (≥ 4) future climate scenarios. Figure was created in R version 3.3.3 [45] (https://www.R-project.org/).

When novel environments are excluded from maps, 31.5% of Australia is projected to be currently suitable for at least one of the species, decreasing to 28% (∼2,100,500 km^2^) by 2070 **(S5 Table**). Hence, exclusion of novel environments substantially impacts the size of suitable habitat (i.e., a substantial area of suitable habitat is projected to occur in areas with novel climatic conditions). However, parts of the Wet Tropics bioregion is projected to remain suitable for all 11 species (70 km^2^) and the major commercial host plants within this bioregion may continue to be at risk of invasion by most or all of these high priority species.

Major commercial host plant regions along the coastal strip of south-east Queensland and north-east NSW are likely to have areas that are suitable under all future scenarios for *B. bryoniae, B. jarvisi, C. capitata* and *Z. cucumis* **(S2, S5, S10 and S11 Figs).** Under some scenarios, these regions may also be suitable for *B. halfordiae, B. neohumeralis* and *B. tryoni* **(S4, S8 and S9 Figs)**. Some major commercial host plant regions in southern NSW and Victoria are also projected to be suitable for *B. jarvisi, B. tryoni* and *C. capitata* under all scenarios **(S5, S9 and S10 Figs)** and for *B. halfordiae, B. neohumeralis* and *Z. cucumis* under a limited number of scenarios **(S4, S8 and S11 Figs)**.

In south-west Western Australia, major horticulture regions are likely to remain suitable for *B. jarvisi, B. tryoni* and *C. capitata*, although the latter species is currently not found in this region **(S5, S9 and S10 Figs).** Commercial horticulture regions in the top-end of the Northern Territory are also likely to be suitable for *B. jarvisi, B. kraussi, B. musae, B. tryoni* and *Z. cucumis* under all scenarios, and *B. frauenfeldi* under some climate scenarios. Horticultural regions in Tasmania are projected as suitable for *B. jarvisi, B. tryoni* and *C. capitata* **(S5, S9 and S10 Figs).**

## Discussion

Our study suggests that the Wet Tropics bioregion has climatically suitable habitat for the largest number of high priority tephritid pest species both now and as a result of climate changes projected to occur through to 2070. Cape York Peninsula and the Northern Territory are also likely to be vulnerable, although novel climates are projected to occur in these regions, and the extrapolation of SDMs to these conditions may be unreliable. The east coast of Australia is also likely to remain suitable for multiple species until at least 2070. As such, major horticulture regions in north-western Australia, the Northern Territory, southern-central regions of NSW, southern Victoria and north Tasmania may become increasingly suitable to high priority fruit flies. Two species, *B. tryoni* (Qfly) and *C. capitata* (Medfly), are projected to have suitable conditions in all states and territories of Australia, under all considered climate change scenarios, until at least 2070.

Models for both Qfly and Medfly were driven primarily by temperature parameters, rather than precipitation. Previous studies have identified climatic constraints on the distribution of Qfly. For instance, it has been reported that Qfly pupae do not survive in the winter months in Melbourne and near Sydney [56], and adults fail to emerge later than mid-April [57]. Further, many subtropical sites in Queensland are marginal in winter for Qfly breeding and general activity [8, 57]. As such, slight temperature increases associated with climate change are projected to substantially elevate the threat that this species poses to horticultural industries [58]. For instance, using data from the late 1990s, it was estimated that annual control costs for apple growers around Adelaide may increase by between $346,000 and $1.3 million with a 0.5–2°C increase in temperature [58].

With the exception of Western Australia, all Australian states and territories are currently free from Medfly, with market access protocols inhibiting movement into other states [21], and incursions met with immediate eradication programs [13]. Our model of current habitat indicates suitable conditions for Medfly around most of Australia’s coastal regions. In addition to identifying suitability in the subtropical coastal fringe of Queensland, our model suggested that much of the low-altitude regions in the south-east, including parts of Tasmania, are also suitable. This is consistent with previous work using CLIMEX to estimate the potential distribution of Medfly [25]. Competition with Qfly may be responsible for exclusion of Medfly from much of Queensland [25], and similar biotic interactions may suppress the species elsewhere [13]. However, Medfly may be more tolerant to low temperatures and dry summers than Qfly [4], rendering Medfly the stronger competitor in areas with these conditions. Medfly was recorded in Tasmania in the 1920s but reportedly failed to survive an unseasonably hot and dry summer [4]. Due to their age, these records were not used to calibrate our model, yet our projections indicate that Tasmania continues to have conditions suitable for this species. Indeed, while strong market access protocols have excluded Qfly from Tasmania for around 90 years, our projections of Tasmania’s suitability were validated upon discovery of an outbreak in 2018.

Allwood and Angeles [59] reported that *B. jarvisi* is recognized as a pest in north-western Australia, infesting mango, guava and pomegranates (as reported in [60]). Dominiak and Worsley [9] concluded that the current south-eastern range limit lies north of the Queensland-NSW border (∼25.5° south), while the south-western limit lies at approximately 18° south.

However, previous analysis suggested that this species’ current climatic range could extend into the cooler temperate areas of southern NSW, and eastern and northern Victoria [4]. Our models partly agree, indicating that suitable conditions currently occur along the east coast of Victoria. This species can also withstand very warm conditions, with eggs known to be more heat tolerant than those of the sympatric Qfly, surviving temperatures of 48.2°C [60]. Given that these species infest many of the same hosts, competition is likely, hence eradication of Qfly may result in the competitive release of *B. jarvisi*, increasing the threat it poses to horticulture [4, 60]. Further, as the cultivation of *B. jarvisi* host plants expands geographically, this species may increase in abundance and extend its range, potentially becoming a major pest in north-western Australia [6, 36].

While widespread throughout Queensland, *Z. cucumis* currently has a restricted distribution in the Northern Territory, although there is a disputed single record from northern Western Australia [61]. Both Fitt [62] and the Horticultural Policy Council [4] reported that if the cucurbit industry expands in the Northern Territory, the pest status of *Z. cucumis* may increase. However, while the species has been trapped frequently in the Northern Territory, it has not been found on cucurbits growing in this region [6]. In NSW, *Z. cucumis* appears to be currently limited to regions close to the Queensland border, with rare detection as far south as Sydney [61]. It has not been detected in the (former) Fruit Fly Exclusion Zone in southern NSW [37]. Our model also estimates the southern limit of suitable climate for this species to be around Sydney. However, with climate change this may extend further southward, with parts of Victoria projected to become increasingly suitable over time, depending on the climate change scenario.

*Bactrocera neohumeralis* presently occurs from the western Cape York Peninsula, Queensland, south to Sydney, NSW [3, 7, 37]. Our models agree that suitable conditions occur from Cape York Peninsula to north of Sydney, although as climate changes, the range of this species may extend southward and, under some scenarios, into parts of Victoria. Previous climatic analysis also suggested that this species is well adapted to conditions on the east coast of Queensland, with large populations occurring in areas north of Townsville [4]. Similar ecological characteristics are shared by *B. neohumeralis* and Qfly [63], yet while Qfly is prevalent in sub-tropical and temperate areas of Queensland and NSW, *B. neohumeralis* is more prevalent in northern wet tropical areas [4, 5, 64]. The reason for this difference between the geographical ranges of these species is unclear, as both are polyphagous and use similar host fruits for their larval development [63, 64].

Our model for *B. aquilonis* indicates that suitable conditions for this species are currently found in northern Queensland, although it is presently only known from north-western Australia [5]. The hosts of this species now include 40 commercial crops [6]. Expansion of the range of this species, or the growth of host plant industries in north-western Australia may necessitate the development of new monitoring, control and disinfestation procedures [60]. In addition, it has been argued that if *B. aquilonis* hybridises with Qfly, and the resulting strain may have greater potential for spread than *B. aquilonis* [4]. This, in turn, would require that disinfestation procedures should be developed for the hybrids [60].

The Australian distribution of *B. bryoniae* ranges from the Torres Strait Islands, across northern Australia, and along the east coast to north of Sydney, NSW. Our results indicate that suitable habitat exists in Victoria, i.e. south of the species’ known range. However, previous studies have demonstrated that populations in northern NSW experience a marked decline in abundance through November–January [37]. This may be explained by a decline in the fruiting and flowering of native host trees, or seasonal climatic constraints that are not reflected in our model [37], which may also explain their absence in Victoria.

Northern Queensland has the highest diversity of fruit flies in Australia, and some species with significant economic impacts are found only in this region [7]. The distribution of *B. kraussi, B. musae* and *B. frauenfeldi* is limited to north Queensland [3, 65], with recent trap data suggesting that these species do not occur south of Townsville [7]. Royer et al. [35] predicted that *B. frauenfeldi* also has suitable habitat in the Northern Territory and northern Western Australia, which is also suggested by our model. The availability of hosts does not appear to limit the range of *B. frauenfeldi*, which has expanded in northern Queensland due to continued planting of hosts, such as mango and guava [35]. Further increases within these horticulture industries in northern Queensland may increase the pest status of this fly [65].

### Model Errors and Uncertainties

SDMs are useful for developing a broad understanding of how the distribution of suitable habitat may be influenced by climate change. However, the output of SDMs is known to be influenced by characteristics of the occurrence sample, including its size [66], sampling bias [67], and spatial autocorrelation [68], as well as the extent of the study area, selection of predictor variables [69], and selection of background points [70]. We addressed these issues by: (1) exploring alternate settings in Maxent to optimise models and reduce overfitting that may generate unreliable estimates [47]; (3) reducing the number of predictor variables by assessing collinearity; and (4) critically examining response curves.

In addition, we acknowledge that the selection of a threshold for converting Maxent’s continuous output into binary data (typically defined as distinguishing between “suitable” and “unsuitable” conditions) can be subjective and problematic. A region classified as unsuitable may not be free of the pest; rather, these areas are considered less likely to support a population compared with regions above the threshold. In reality, the choice of threshold is based upon a comparison of the importance of false positives (FP) and false negatives (FN) [71]. For invasive species, the latter may be more serious because they can result in an underestimate of the geographic extent of suitable habitat, and hence, invasion risk [72]. This, in turn, can lead to poor decision-making and failure to establish appropriate surveillance or containment measures. Hence, in this context a precautionary approach to defining a threshold, as undertaken in the present study, is warranted. However, since overprediction of suitable habitat can also prove problematic (potentially leading to ineffective allocation of monitoring resources), we provide continuous (unthresholded) model output, permitting stakeholders to modify this threshold according to their objectives.

Sampling bias is another challenge faced when fitting correlative SDMs, particularly when incorporating data from sources of incidental observations such as museums and natural history collections [73]. As such, it is difficult to determine whether a species is observed in a particular environment because of habitat preferences or because that region has received the largest search effort [70, 73]. For presence-background approaches to habitat modelling, a target-group background sampling strategy goes some way to handling biased occurrence samples [74]. However, while imposing environmental bias on the background counteracts similar bias in the occurrence sample, this strategy may increase the extent of novel environments to which the model must be extrapolated.

While SDMs consider exposure to climate change, species responses may also include microevolution [75] or plasticity [76]. As accessibility to genomic data increases, and experiments on plasticity are conducted, SDM output can be refined [77]. In addition, as mean conditions change, so too will the distribution and magnitude of extremes. Presently, there has been little work undertaken to assess how different fruit fly species tolerate extreme weather events such as heatwaves and moisture stress.

We also note that our analysis does not take into consideration the potential necessity for horticultural industries to shift geographically to adapt to climate change. Analysing potential shifts in climatic suitability for horticultural crops is complicated by our capacity to modify the environment (e.g. through irrigation), and thus was beyond the scope of this study.

## Conclusions

Surveillance activities, pre- and post-harvest treatment, and control activities for fruit flies present a substantial cost to Australia’s horticultural industries [2, 4, 14]. Climate change is likely to alter the distribution of suitable habitat for these species, and this could have ramifications for Australia’s horticultural industries. Our analysis highlights that the major horticultural production regions are likely to remain suitable for multiple economically important fruit fly species as climate changes. Furthermore, given that knowledge of current species distributions remains the basis for market access decisions, the potential for range shifts to occur is of critical interest to horticultural industries.

Our model projections identify geographic regions likely to be suitable for 11 high priority, economically important pest fruit fly species, now and in the future. Outputs from this study provide guidance to pest managers, such that they can assess pest risks and design appropriate ongoing surveillance strategies. Our results emphasize the importance of vigilance and preparedness across Australia, to prevent further range expansion of these 11 species, and underscore the need for ongoing research and development into monitoring, control, and eradication tools.

## Supporting Information

**S1-11 Figs. Climatic habitat suitability for 11 tephritid fruit flies under various future climate scenarios.** (1) *Bactrocera aquilonis* (2) *Bactrocera bryoniae* (3) *Bactrocera frauenfeldi* (4) *Bactrocera halfordiae* (5) *Bactrocera jarvisi* (6) *Bactrocera kraussi* (7) *Bactrocera musae* (8) *Bactrocera neohumeralis* (9) *Bactrocera tryoni* (10) *Ceratitis capitata.* (11) *Zeugodacus cucumis*. (A) current habitat suitability modelled using Maxent – values close to zero represent areas with low climatic suitability while values closer to one indicate higher climatic suitability; (B) areas considered “suitable” (i.e., with habitat suitability values above the 10th percentile at training presence sites, shown in red); (C, D, E) agreement about the suitability of habitat for the species across six climate scenarios for 2030, 2050 and 2070, respectively; (F) the location of Australian occurrence records of the species, based on specimens from natural history collections, literature and State Government trapping programs, and major commercial horticultural hosts, according to the Australian Horticulture Statistics Handbook (HSHB; www.horticulture.com.au).

**S12 Fig. Hotspot maps of habitat suitability for the 11 fruit fly species under climate change, when novel environments are excluded.** Hotspot maps of current and future habitat suitability for 11 fruit flies with regions containing novel environments considered unsuitable. Suitability was modelled with Maxent, and thresholded using the 10th percentile at training presence localities. Colours indicate the number of species for which habitat is predicted to be suitable under the majority (≥ 4) future climate scenarios. Figure was created in R version 3.3.3[1] (https://www.R-project.org/).

**S1 Table. Model performance and bioclimatic variables used to investigate the suitability of habitat for tephritid fruit fly species.** AUC value indicates the area under the receiver operating characteristic curve (average of 5 cross-validated replicates), which was used to evaluate model performance; SD (standard deviation); and HPI (highest permutation importance, %) of bioclimatic variables contributing to the model where BIO01: annual mean temperature, BIO02: mean diurnal range; BIO03: isothermality; BIO06: minimum temperature of the coldest month; BIO07: temperature annual range; BIO11: mean temperature of the coldest quarter; BIO13: precipitation of wettest month; BIO14: precipitation of the driest month; BIO16: precipitation of the wettest quarter; BIO17: precipitation of the driest quarter and BIO19: precipitation of the coldest quarter.

**S2 Table. Projected changes in the area of suitable habitat for all 11 fruit fly species, under six future climate scenarios, relative to the current period.** (1) *Bactrocera aquilonis* (2) *Bactrocera bryoniae* (3) *Bactrocera frauenfeldi* (4) *Bactrocera halfordiae* (5) *Bactrocera jarvisi* (6) *Bactrocera kraussi* (7) *Bactrocera musae* (8) *Bactrocera neohumeralis* (9) *Bactrocera tryoni* (10) *Ceratitis capitata* (11) *Zeugodacus cucumis*. For each species, the first column indicates the GCM (Global Climate Model) for three time periods 2030, 2050 and 2070. Other columns: % Lost refers to the percentage of currently suitable habitat projected to become unsuitable in the future; % Gained refers to the percentage of future suitable habitat that is in areas currently unsuitable; Range Changed refers to the change (%) between the size of current and future suitable habitat (positive numbers indicate an increase in range size, negative numbers indicate a decrease).

**S3 Table. Area (km**^2^**) and percentage of Australia projected to be suitable for 11 fruit flies under six future climate scenarios.** In the column ‘Climate scenarios’, 0 refers to the area projected to be unsuitable across all six scenarios; 1 refers to the area projected to be suitable under any one of the six scenarios…6 refers to the area projected to be suitable under all six scenarios.

**S4 Table. Major commercial fruits and vegetables host species to the Australian Horticulture Statistics Handbook (HSHB;** www.horticulture.com.au). Pest status is based on Hancock et al^3^, where “major” indicates that there have been many records of the fly infesting that host.

**S5 Table. Area (km**^2^**) and percentage (%) of Australia projected to be suitable for the 11 fruit fly species considered in this study, when novel environments have been excluded.** Each row of the table indicates the area (and percentage) projected to be suitable now, in 2030, 2050, and 2070, for *n* species, where *n* is given in the “Count” column. Thus, the first row (with Count = 0) gives the area projected to be unsuitable for all 11 species, the row with Count = 1 gives the area projected to be suitable for any one of the 11 species, and the row with Count = 11 gives the area projected to be suitable for all 11 species. Values shown here apply when cells with novel environmental conditions are considered unsuitable.

### Acknowledgements

We gratefully acknowledge our data providers Nick Secomb (Plant Health Operations Biosecurity, PIRSA, South Australia) and Lauren Donaldson (Department of Economic Development, Jobs, Transport and Resources, Victoria). Special thanks to Phil Taylor, Dan Ryan, and Penny Measham for their feedback and advice. SS was supported by an International Macquarie University Research Excellence Scholarship (iMQRES). This research was conducted as part of the SITplus collaborative fruit fly program.

## Author contributors

Data were collated by SS, JBB, BCD, JER, and LJB. Models were calibrated by SS and evaluated by JBB, BCD, JER, and LJB. Manuscript was drafted by SS with contributions from JBB, BCD, JER, and LJB.

